# Functional degradation: a mechanism of NLRP1 inflammasome activation by diverse pathogen enzymes

**DOI:** 10.1101/317834

**Authors:** Andrew Sandstrom, Patrick S. Mitchell, Lisa Goers, Edward W. Mu, Cammie F. Lesser, Russell E. Vance

**Author notes:** These authors contributed equally to this work (Andrew Sandstrom, Patrick S. Mitchell).

## Abstract

Inflammasomes are multi-protein platforms that initiate innate immunity by recruitment and activation of Caspase-1. The NLRP1B inflammasome is activated upon direct cleavage by the anthrax lethal toxin protease. However, the mechanism by which cleavage results in NLRP1B activation is unknown. Here we find that cleavage results in proteasome-mediated degradation of the N-terminal domains of NLRP1B, liberating a C-terminal fragment that is a potent Caspase-1 activator. Proteasome-mediated degradation of NLRP1B is both necessary and sufficient for NLRP1B activation. Consistent with our new ‘functional degradation’ model, we identify IpaH7.8, a *Shigella flexneri* ubiquitin ligase secreted effector, as an enzyme that induces NLRP1B degradation and activation. Our results provide a unified mechanism for NLRP1B activation by diverse pathogen-encoded enzymatic activities.

**One Sentence Summary:** Two distinct pathogen enzymes activate an innate immune sensor called NLRP1B by a mechanism that requires proteasome-mediated degradation of NLRP1B.

## Main Text

In animals, pathogens are generally recognized by germline-encoded innate immune receptors that bind directly to conserved pathogen-associated molecular patterns (PAMPs) such as bacterial lipopolysaccharide or flagellin (*1*). Recognition of PAMPs permits robust self-nonself discrimination, but because PAMPs are found on harmless as well as pathogenic microbes, PAMP receptors do not distinguish pathogens from non-pathogens. Plants also use germline-encoded receptors to detect PAMPs, but in addition, respond to infection by indirect detection of secreted pathogen enzymes called ‘effectors’ (*2*). In this mode of recognition, called ‘effector-triggered immunity’, intracellular proteins of the nucleotide-binding domain leucine-rich repeat (NLR) superfamily sense effector-induced perturbation of host signaling pathways. Because harmless microbes do not deliver effectors into host cells, effector-triggered immunity is inherently pathogen-specific. It has been proposed that animals may also detect pathogen-encoded activities (*3-10*), but there are relatively few examples of this mode of pathogen recognition that are understood in molecular detail.

We sought to determine how a particular mammalian NLR protein called NLRP1 senses pathogen-encoded activities. NLRP1 is the founding member of a class of proteins that form inflammasomes (*11*), multi-protein platforms that initiate immune responses by recruiting and activating pro-inflammatory proteases, including Caspase-1 (CASP1) (*12-14*). CASP1 cleaves and activates specific cytokines (interleukins-1β and −18) and a pore-forming protein called Gasdermin D, leading to a host cell death called pyroptosis. In certain strains of mice and rats, a specific NLRP1 family member called NLRP1B is activated via direct proteolysis of its N-terminus by the lethal factor (LF) protease secreted by *Bacillus anthracis* (*15-18*). Previous studies demonstrated that N-terminal proteolysis is sufficient to initiate NLRP1B inflammasome activation (*16*), but the molecular mechanism by which proteolysis activates NLRP1B has been elusive.

Similar to other NLRs, NLRP1B contains a nucleotide-binding domain and leucine-rich repeats (Fig. 1A). However, NLRP1B also exhibits several unique features. First, the NLRP1B caspase activation and recruitment domain (CARD) is C-terminal instead of N-terminal, as in other NLRs. Second, NLRP1B is the only NLR that contains a function-to-find domain (FIIND). The FIIND constitutively undergoes an auto-proteolytic event, resulting in two separate NLRP1B polypeptides that remain non-covalently associated with each other (*19-21*). Mutations that abolish FIIND auto-processing block inflammasome activation (*20, 21*), but it remains unclear why FIIND auto-processing is essential for NLRP1B function. Lastly, proteasome inhibitors specifically block NLRP1B inflammasome activation, but do not affect other inflammasomes or inhibit LF protease, suggesting the proteasome is specifically required for activation of NLRP1B itself (*22-26*).

**Fig. 1.**
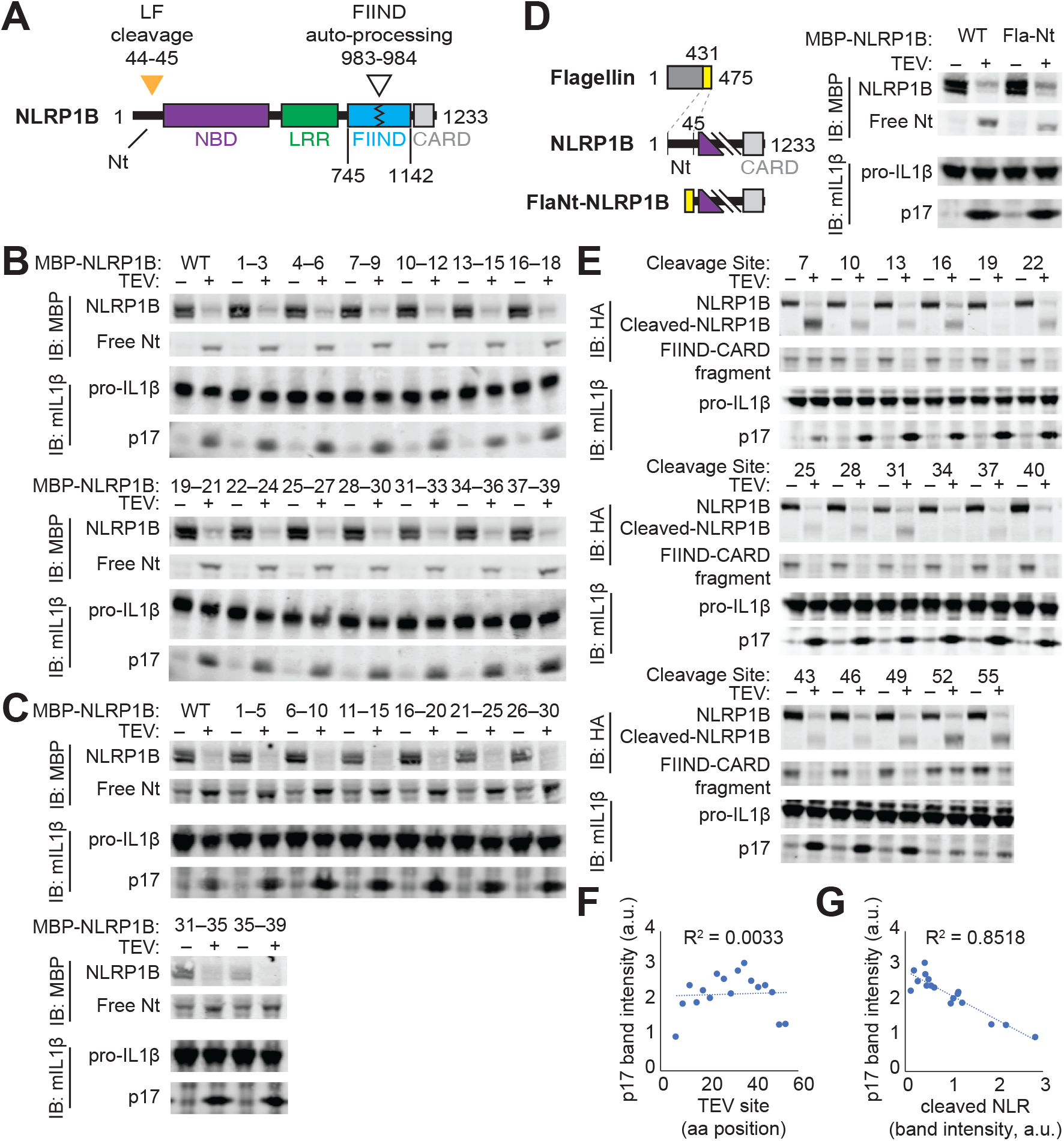
The N-terminal domain of NLRP1B does not mediate auto-inhibition. (**A**) Schematic of mouse NLRP1B domain architecture. Nt, N-terminus; NBD, nucleotide-binding domain; LRR, leucine-rich repeat; FIIND, function-to-find domain; CARD, caspase activation and recruitment domain. FIIND auto-processing is usually not complete and thus NLRP1 appears as a doublet [(B to D), upper blot] when probing with an antibody directed against the N-terminal MBP tag. The location of lethal factor (LF) cleavage (orange triangle) and FIIND auto-processing (white triangle) is shown. (**B**, **C**) Mutagenesis of the Nt does not disrupt auto-inhibition. The effect of sequential, non-overlapping replacements of Nt amino acids 1-39 by triple alanine [(B), AAA] or glycine-serine-glycine [(C), GGSGG] scanning mutagenesis on NLRP1B auto-inhibition was assessed. In (B to E) inflammasome activation was induced by co-expression of the tobacco-etch virus (TEV) protease which cleaves a TEV site engineered into MBP-NLRP1B. Activation was monitored by CASP1 processing of pro-IL-1β to p17. (**D**) Replacement of the Nt of NLRP1B with a heterologous sequence from flagellin (FlaNt) does not affect auto-inhibition or protease-dependent activation of NLRP1B. (**E** to **G**) NLRP1B degradation, but not the position of protease cleavage, positively correlates with IL-1β processing. The TEV site was positioned sequentially along the NLRP1B Nt and TEV cleavage was detected by probing for a C-terminal HA tag [(E), upper blot]. IL-1β processing was plotted relative to the position of the TEV site (F) or the protein level of cleaved-NLRP1B (G). IB, immunoblot.

Cleavage of NLRP1B by LF results in a loss of 44 amino acids from the N-terminus of NLRP1B, an event that correlates with its activation (*15-18*). We and others have previously proposed an ‘auto-inhibition’ model to explain NLRP1B activation (*13, 27*). In this model, the N-terminus of NLRP1B functions as an auto-inhibitory domain that is lost after cleavage by LF. The NLRP1B N-terminus might mediate auto-inhibition either through direct engagement with other NLRP1B domains in *cis*, or by binding to an inhibitory co-factor. A clear prediction of the auto-inhibition model is that sequences within the N-terminus should be required to prevent spontaneous inflammasome activation. To identify such sequences, we systematically mutated the NLRP1B N-terminus by replacing groups of three consecutive amino acids with alanines (Fig. 1B) or by replacing groups of five sequential amino acids with a flexible GGSGG motif (Fig. 1C). Each mutant was also engineered to contain an N-terminal TEV protease site to enable inducible NLRP1B cleavage and activation (*16, 28*). Inflammasome activity was monitored by CASP1-dependent processing of pro-IL-1β to p17 in a reconstituted inflammasome system in transfected 293T cells (*16, 28*). To our surprise, none of the mutants was auto-active, even though all showed full activity upon cleavage. To test if any N-terminal sequence could mediate auto-inhibition, we replaced the entire N-terminus with a heterologous alpha-helical domain from bacterial flagellin. Again, to our surprise, the hybrid Fla-NLRP1B protein was not auto-active, but was still functional after N-terminal cleavage (Fig. 1D). Together with prior experiments (*20, 28, 29*), these results caused us to reconsider a model in which specific N-terminal sequences of NLRP1B mediate auto-inhibition.

We then asked whether the precise site of N-terminal proteolysis is a major determinant of NLRP1B activation. We generated a series of NLRP1B variants in which a TEV protease cleavage site was positioned at regular intervals from the N-terminus. We found that cleavage of as few as 10 amino acids from the N-terminus was sufficient to activate NLRP1B. Furthermore, there was not a significant correlation between the position of TEV cleavage and NLRP1B activity (Fig. 1E, F). In contrast, we noticed a striking negative correlation between the amount of TEV-cleaved NLRP1B protein and inflammasome activation (Fig. 1E, G). This correlation, as well as prior evidence that proteasome inhibitors block NLRP1B activity (*22-26*), led us to consider whether proteasome-mediated degradation of NLRP1B is an important step in its activation. Consistent with this idea, we found that the proteasome inhibitors MG132 and Bortezomib not only abrogated NLRP1B activation in our reconstituted 293T system, but also prevented loss of the cleaved NLRP1B protein after LF treatment (Fig. 2A). By contrast, inhibition of p97/VCP by NMS-873 had no effect on NLRP1B activation.

**Fig. 2.**
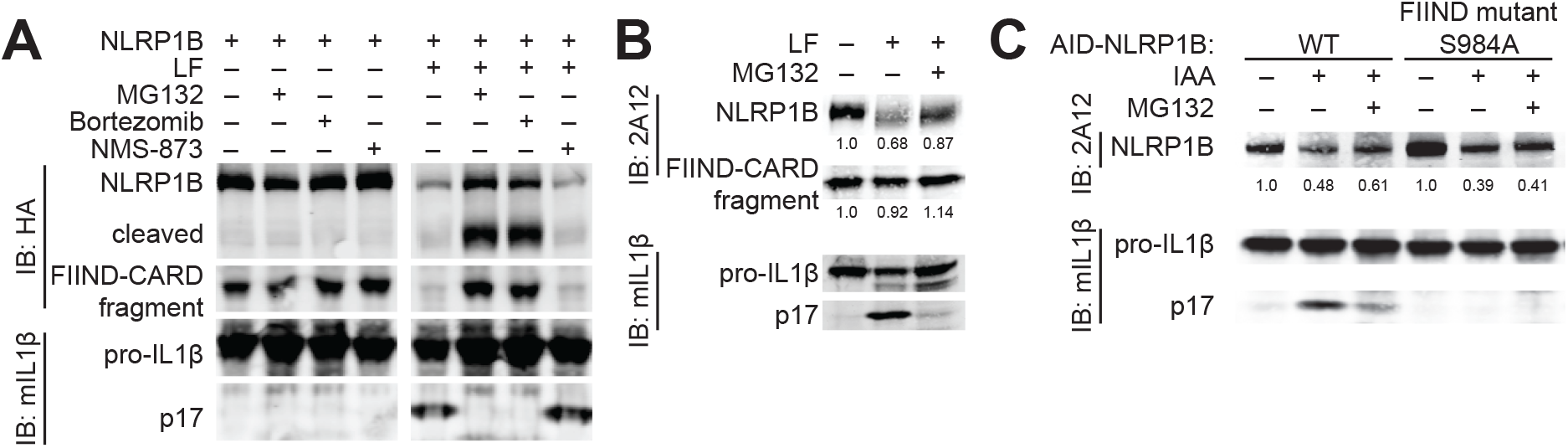
Degradation of NLRP1B is necessary and sufficient for NLRP1B inflammasome activation. (**A** to **B**) The proteasome is required for NLRP1B inflammasome activation. Proteasome inhibitors MG132 (10μM) and Bortezomib (1μM) block LF-induced NLRP1B degradation and IL-1β processing in transfected 293T cells (A) as in Fig. 1 and immortalized 129 bone-marrow-derived macrophages (B). Inhibition of p97/VCP by NMS-873 (0.5μM) had no effect on IL-1β processing in transfected 293T cells. (**C**) Proteasomal degradation of NLRP1B is sufficient for inflammasome activation. The plant auxin-interacting degron (AID) was fused to the N-terminus of indicated GFP-NLRP1B variants; specific degradation was induced with Indole-3-acetic acid (IAA) in TIR1-expressing 293T cells. IAA-induced NLRP1B degradation induced IL-1β processing, which is dependent on FIIND auto-processing and is blocked by proteasome inhibitors. The average fold change (FC) of the NLRP1B band signal intensity in LF, IAA or IAA+MG132 treated versus untreated samples is indicated below the NLRP1B blot (B and C). S, serine. A, alanine.

To determine if endogenous NLRP1 is also lost from cells after cleavage, we derived a monoclonal antibody (2A12) against the C-terminal CARD domain of NLRP1B (*Fig. S1*). Using this antibody, we could track endogenous NLRP1B in 129S1/SvimJ (129) bone marrow-derived macrophages (BMMs) after treatment with LF (Fig. 2B). As observed in the 293T system, LF treatment led to a loss of full-length NLRP1B, which was at least partially reversed by MG132 treatment. In contrast, proteasome inhibitors have no effect on NLRP3 (*23, 26*) or NAIP5/NLRC4 inflammasome activation (*Fig. S2*), consistent with the hypothesis that the proteasome functions uniquely in NLRP1B inflammasome activation.

The precise mechanism by which NLRP1B is targeted to the proteasome after LF cleavage remains unknown. However, the N-end rule pathway is known to recognize cleaved proteins (*30, 31*), and N-end rule inhibitors block NLRP1B activation (*25*). Therefore, we propose that cleavage of NLRP1B reveals a destabilizing neo-N-terminus that targets NLRP1B for ubiquitylation by N-end rule E3 ubiquitin ligases. Consistent with this hypothesis, in independent and parallel work, the Bachovchin laboratory has recently identified N-end rule ubiquitin ligases that are critical for LF-mediated NLRP1B activation [Chui, A.J. *et al* (2018) *Biorxiv.org*]. We note that all the TEV-cleavable NLRP1B variants we examined generate a glycine as the neo-N-terminus after cleavage, which the N-end rule predicts to be stabilizing (*32*) despite our observation that TEV-cleavage results in NLRP1B destabilization (Fig. 1).

Importantly, however, previous work has shown that aminopeptidase inhibitors are able to block LF-mediated killing of RAW264.7 cells (*25*). Thus, it appears likely that after an initial cleavage event, aminopeptidases act to remove N-terminal amino acids and expose internal amino acids to N-end rule recognition. Consistent with this idea, we swapped the P2′ residues between two differentially activated TEV-cleavable NLRP1B variants and found that the P2′ residue can also modulate activity (*Fig. S3*). Thus, the stability and activity of cleaved NLRP1B depends on more than just the identity of the neo-N-terminal amino acid, consistent with a growing body of evidence that multiple determinants (e.g., N-acetylation) underlie N-end rule degradation (*33*).

In the above experiments, the inhibition of NLRP1B activation by proteasome inhibitors might have been due to stabilization of a negative regulator of NLRP1B rather than to stabilization of NLRP1B itself. Therefore, we next asked whether specific degradation of NLRP1B is sufficient to induce its activation. To achieve selective degradation of NLRP1B, the auxin-interacting degron (AID) (*34, 35*) was fused to the N-terminus of NLRP1B. Upon addition of the auxin hormone indole-3-acetic acid (IAA), the AID recruits a co-expressed TIR1 E3 ligase that specifically ubiquitylates AID-fusion proteins, targeting them to the proteasome. Indeed, IAA induced rapid degradation of the AID-NLRP1B fusion protein and stimulated robust IL-1β processing (Fig. 2C). Of note, we found that FIIND auto-processing is also required for IAA-induced activation of AID-NLRP1B (Fig. 2C). Together these results demonstrate that proteasomal degradation of NLRP1B itself is both necessary and sufficient for activation of the NLRP1B inflammasome.

To explain the seemingly paradoxical observation that NLRP1B degradation leads to its activation, we propose the following ‘functional degradation’ model (Fig. 3A). Our proposal relies on the prior observation that FIIND auto-processing is required for activation of NLRP1B. The FIIND domain comprises two separate sub-domains, termed ZU5 and UPA, with auto-processing occurring near the C-terminal end of the ZU5 domain (at F983|S984) (*19-21*). After auto-processing, the C-terminal fragment of NLRP1B thus consists of the UPA domain fused to the CARD that is required for CASP1 recruitment and activation. Prior to activation, the FIIND(UPA)-CARD fragment is non-covalently associated with the rest of NLRP1B (Fig. 3A). After cleavage by LF, NLRP1B is targeted to the proteasome, a processive protease that degrades polypeptides by feeding them through a central barrel (*36*). Critically, however, directional (N-to C-terminus) and processive degradation of NLRP1B by the proteasome will be interrupted by the covalent break within the auto-processed FIIND domain. At this point, we propose that the C-terminal FIIND(UPA)-CARD fragment is released and is able to seed inflammasome assembly (Fig. 3A).

**Fig. 3.**
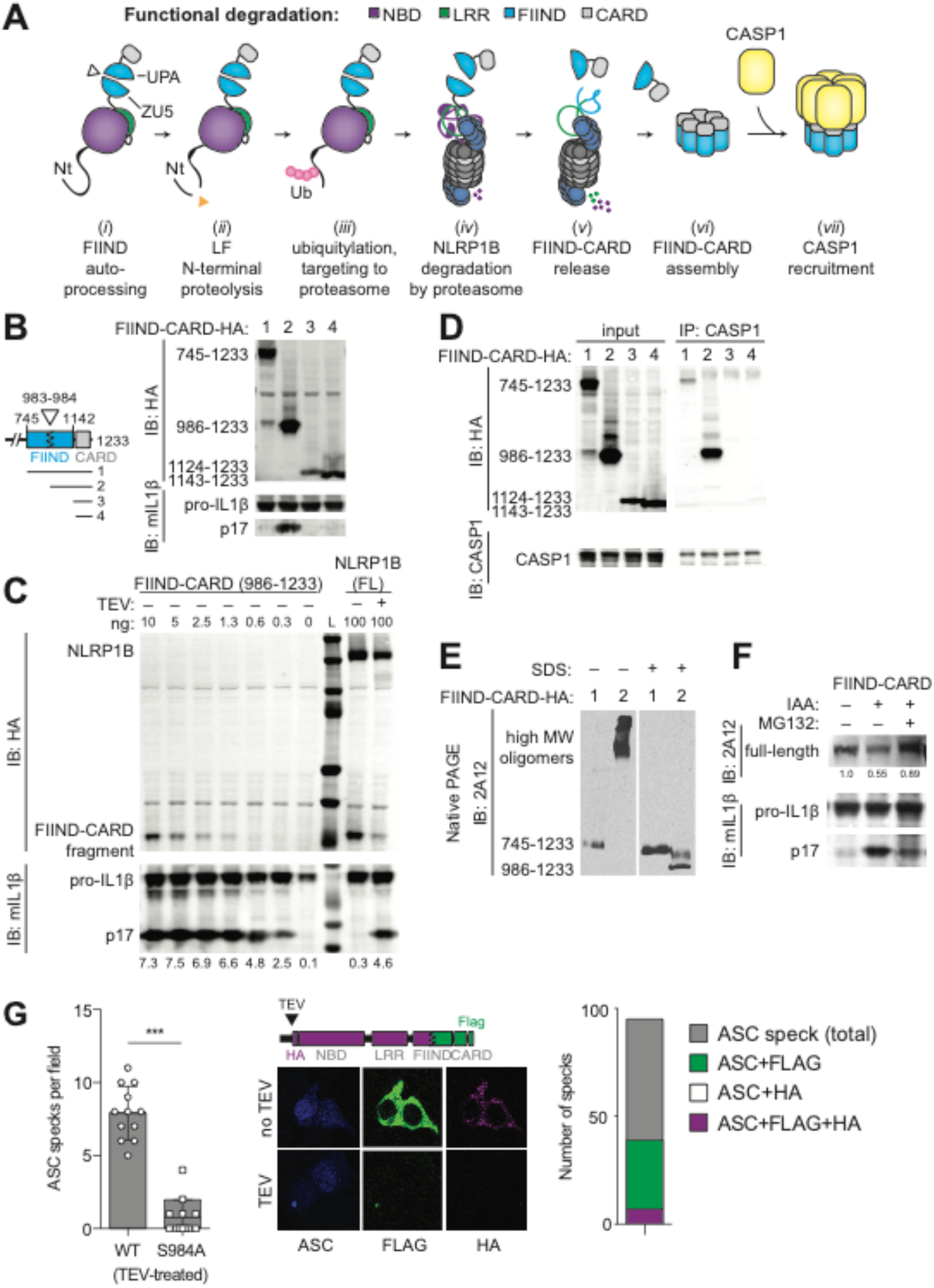
‘Functional degradation’ of NLRP1B liberates the FIIND(UPA)-CARD fragment, a highly potent inflammasome activator. (**A**) A model for NLRP1B activation via ‘functional degradation’: (*i*) constitutive auto-processing of the NLRP1B FIIND domain results in two non-covalently associated polypeptides: NBD-LRR-FIIND(ZU5) and FIIND(UPA)-CARD; (*ii*) Lethal factor (LF) protease cleavage of the NLRP1B Nt exposes a neo-Nt; (*iii*) N-end rule factor recognition of the neo-Nt results in ubiquitylation of NLRP1B; (*iv*) NLRP1B is degraded by the proteasome, resulting in (*v*) release of the FIIND(UPA)-CARD fragment; (*vi*) The FIIND(UPA)- CARD fragment self-assembles into a high molecular weight oligomer which (*vii*) serves as a platform for CASP1 maturation and downstream inflammasome signaling. (**B**) The FIIND(UPA)-CARD fragment has inflammasome activity. The activity of the indicated C-terminal HA-tagged variants was tested in 293T cells as in Fig. 1. Only the FIIND(UPA)-CARD fragment exhibited auto-activity. (**C**) The FIIND(UPA)-CARD fragment is a potent activator of IL-1β processing. Decreasing amounts (ng) of plasmid encoding the FIIND(UPA)-CARD fragment (left) compared to full-length NLRP1B activated by TEV (right). Signal intensity of the p17 band is indicated below the blot. L, Protein ladder. (**D**) The FIIND(UPA)-CARD fragment, but not the full length FIIND(ZU5+UPA)-CARD, truncated FIIND(UPA)-CARD fragment or the CARD domain only, co-immunoprecipitates with CASP1. (**E**) The FIIND(UPA)-CARD fragment, but not the full length FIIND(ZU5+UPA)-CARD forms a high molecular weight oligomer on native PAGE. For (D and E) lanes and FIIND-CARD variants are labeled as in (C). Proteins were native or denatured with sodium dodecyl sulfate (SDS) as indicated. (**F**) IAA-induced degradation of AID-FIIND-CARD is sufficient for inflammasome activation. An AID-FIIND(ZU5+UPA)-CARD is activated by IAA and is blocked by MG132. The average fold change (FC) of the NLRP1B band signal intensity in IAA or IAA+MG132 treated versus untreated samples is indicated below the NLRP1B blot. (**G**) The FIIND(UPA)-CARD fragment co-localizes with the ASC speck. 293T cells were transfected with constructs producing ASC (blue) and an NLRP1B variant marked with a C-terminal FLAG (green) and an N-terminal HA (magenta). As schematized, the HA is inserted directly after the TEV cleavage site, allowing detection of the NBD-LRR-ZU5 while the FLAG tag marks the UPA-CARD. The number of ASC specks per field in TEV-treated samples is greatly reduced in cells transfected with the S984A FIIND auto-processing mutant. Representative images depict cytosolic FLAG and HA signal in untreated samples, with FLAG colocalization with ASC specks and concomitant loss of the HA signal in TEV-expressing cells. The total number of ASC specks, or ASC specks positive for both FLAG and HA (n=7) or only FLAG (n=32) or HA (n=0) in TEV-treated samples is quantified from 12 fields. Significance was determined by t-test ^***^, P<0.001.

Our new ‘functional degradation’ model of NLRP1B inflammasome activation has several virtues. First, the model explains how N-terminal cleavage results in proteasome-dependent NLRP1B activation without a requirement for specific N-terminal ‘auto-inhibitory’ sequences. Second, the model accounts for why the NLRP1B CARD is C-terminal, rather than N-terminal, as only the C-terminus of NLRP1B remains after proteasome-mediated degradation. Lastly, the model explains why FIIND domain auto-processing is required for NLRP1B activity: an unprocessed FIIND mutant would be fully degraded and would not release a C-terminal CARD-containing fragment.

A strong prediction of the ‘functional degradation’ model is that the C-terminal FIIND(UPA)-CARD fragment possesses inflammasome activity. Consistent with prior work (*21, 37*), we found that the FIIND(UPA)-CARD fragment was indeed sufficient to promote robust CASP1 activity in our 293T reconstituted inflammasome assay, whereas the full-length FIIND(ZU5+UPA)-CARD appeared inactive despite auto-processing (Fig. 3B). Likewise, the isolated CARD domain (lacking any portion of the FIIND) also appeared inactive, implying that the FIIND(UPA) domain contributes to inflammasome formation. This pattern was observed across a range of expression levels (*Fig. S4*). Importantly, the FIIND(UPA)-CARD fragment appears to be a highly potent activator of inflammatory signaling. By titrating the amount of expression construct, we found that the FIIND(UPA)-CARD fragment appeared to be up to ~150× more potent than TEV-cleavable full-length NLRP1B (Fig. 3C). These results imply that only a tiny fraction of the total NLRP1B in a cell may need to be degraded to liberate sufficient amounts of the FIIND(UPA)-CARD fragment for robust inflammasome activation. Thus, even if NLRP1B is only degraded a fraction of the time in the ‘productive’ N- to C-terminal direction, this would still likely be sufficient for robust inflammasome activation.

Our ‘functional degradation’ model predicts that the mature/assembled NLRP1B inflammasome consists solely of the FIIND(UPA)-CARD fragment and CASP1. In support of this hypothesis, only the UPA-containing FIIND-CARD fragment co-immunoprecipitated robustly with CASP1 (Fig. 3D) and assembled into higher order oligomers when assessed via non-denaturing PAGE (Fig. 3E). As such, our model predicts that the NBD and LRR domains of NLRP1B are dispensable for inflammasome activation. Indeed, prior studies found that NBD and LRR mutants of NLRP1B are active, or are even constitutively active (*27, 28, 37, 38*), a result that can be explained by the functional degradation model if the mutations destabilize NLRP1B. We further observed that IAA-induced degradation of an AID-FIIND(ZU5+UPA)-CARD fragment is sufficient to induce CASP1 activity (Fig. 3F). This result implies that the FIIND-CARD module is sufficient to impart both auto-inhibition and proteasome-induced activation to NLRP1B. While the role of the NBD and LRRs of NLRP1B remains to be determined, we speculate that these domains may contribute to recognition of pathogen-encoded effectors.

To visualize the simultaneous degradation of the N-terminal domains and the release of the FIIND(UPA)-CARD fragment upon NLRP1B activation, we engineered a variant of NLRP1B with a C-terminal FLAG tag, as well as an HA-tag following the N-terminal TEV cleavage site. In unstimulated cells expressing this variant, we observed both FLAG and HA co-staining in the cytosol (Fig. 3G). In contrast, in cells co-transfected with TEV protease, we observed inflammasome activation as indicated by formation of ‘specks’ containing the ASC adaptor protein. Importantly, we found that the FLAG-tagged UPA-CARD fragment, but not the HA-tagged N-terminal domains, robustly formed puncta co-localized at the ASC speck (Fig. 3G). Moreover, we also observed near complete loss of the HA signal, consistent with N-terminal degradation upon TEV cleavage. The FLAG signal was also lost upon TEV cleavage when FIIND processing was disrupted (*Fig. S5*). These results are consistent with our model that the C-terminal UPA-CARD is released to seed inflammasome formation upon proteolytic cleavage and subsequent proteasomal degradation of the NLRP1B N-terminal domains.

A major implication of our ‘functional degradation’ model is that in addition to detecting pathogen-encoded proteases such as LF, NLRP1B can also potentially sense any enzymatic activity that results in NLRP1B degradation. Several pathogens encode E3 ubiquitin ligases that promote virulence through degradation of host target proteins (*39*). We therefore tested whether the type III secretion system (T3SS)-secreted IpaH family of E3 ubiquitin ligases, encoded by the intracellular bacterial pathogen *Shigella flexneri* (*40-42*), are detected by NLRP1B. Using our reconstituted 293T cell system, we found that IpaH7.8, but not IpaH1.4, 4.5 or 9.8, markedly reduced NLRP1B protein levels and induced IL-1β processing in an NLRP1B-dependent manner (Fig. 4A). We also noted IpaH4.5 reduced IL-1β levels in cells, but we did not pursue this observation. IpaH7.8 selectively activated the 129 but not the C57BL/6 (B6) allele of NLRP1B (Fig. 4B). As expected, FIIND auto-processing was required for IpaH7.8-induced NLRP1B activation (Fig. 4C). Truncation of either the IpaH7.8 LRR or E3 domains, as well as mutation of the catalytic cysteine residue (CA) required for E3 ligase activity, also abolished IpaH7.8- mediated inflammasome activation (Fig. 4D). We observed that IpaH7.8, but not IpaH9.8 or an IpaH7.8 catalytic mutant (IpaH7.8CA), was able to directly ubiquitylate the 129 allele, but not the B6 allele, of NLRP1B (Fig. 4E) in a reconstituted ubiquitylation assay. These results show that IpaH7.8/ubiquitylation-dependent degradation of NLRP1B can result in its activation.

**Fig. 4.**
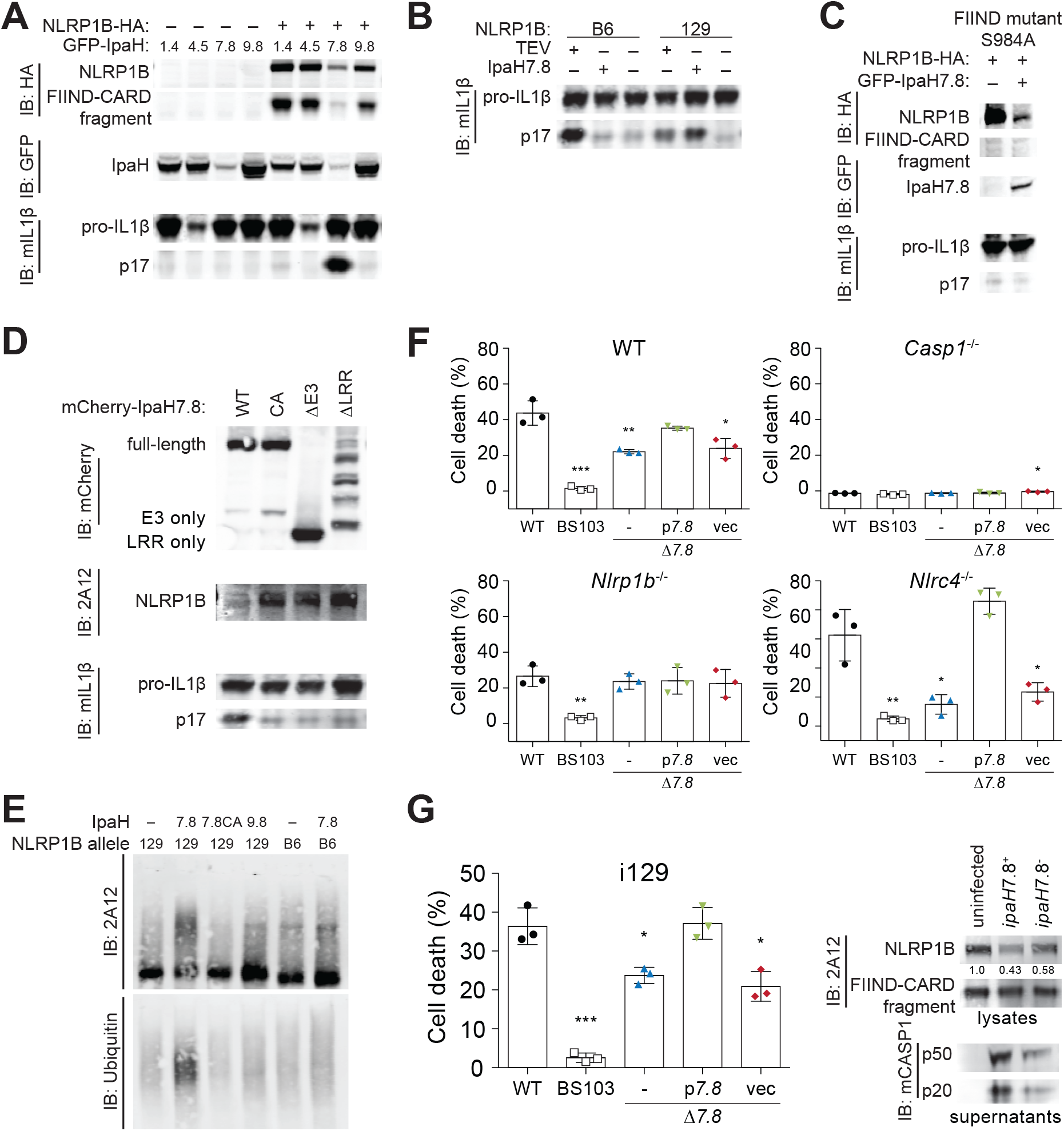
The secreted *Shigella flexneri* IpaH7.8 E3 ubiquitin ligase activates NLRP1B. (**A**) IpaH7.8 induces the degradation and activation of NLRP1B. IpaH7.8, but not IpaH1.4, 4.5 or 9.8, induces NLRP1B degradation and inflammasome activation in 293T cells. (**B**) IpaH7.8 selectively activates the 129 but not the B6 allele of NLRP1B. (**C**) IpaH7.8 induces degradation but not activation of a S984A NLRP1B mutant that is defective for FIIND auto-processing. S, serine. A, alanine. (**D**) Activation of NLRP1B by IpaH7.8 requires essential IpaH7.8 domains. CA, catalytic mutant; ∆E3, deletion of Ub ligase domain; ∆LRR, deletion of LRR. (**E**) IpaH7.8, but not a catalytic mutant (7.8CA) or IpaH9.8, ubiquitylates the 129 allele, but not the B6 allele, of NLRP1B in an in vitro ubiquitylation assay. (**F**) IpaH7.8-mediated macrophage cell death requires NLRP1B. Infection (MOI 30) of WT, *Casp1^−/−^*, *Nlrp1b^−/−^* or *Nlrc4^−/−^* RAW264.7 cells with the WT *Shigella flexneri* strain 2457T (black circle) or mutants strains. BS103, virulence plasmid-cured (white box); Δ*7.8*, *ipaH7.8* deletion (blue triangle); p*7.8*, Δ*7.8* strain complemented with pCMD136 *ipaH7.8* (green inverted triangle); vec, Δ*7.8* strain complemented with pCMD136 empty vector (red diamond). Cell death was monitored by assaying for lactate dehydrogenase (LDH) activity in culture supernatants 30 minutes post-infection. (**G**) IpaH7.8 activates the NLRP1B inflammasome in macrophages. Immortalized 129 (i129) bone-marrow-derived macrophages were infected with *S. flexneri* strains as in (F). IpaH7.8-producing strains induce NLRP1B degradation and CASP1 processing. Cell death was measured by LDH as in (F) 2 hours post-infection. Data sets were analyzed using one-way ANOVA. P values were determined by Dunnet’s multiple comparison post test. ^*^, P < 0.05; ^**^ P < 0.01; ^***^P < 0.001.

*S. flexneri* robustly activates multiple inflammasomes (*43*) and has previously been reported to cause macrophage cell death in an IpaH7.8- and NLRP1B-dependent manner (*44-46*), but a connection between IpaH7.8 and NLRP1B has not been established. Consistent with prior studies (*44, 46*), we found that wild-type *S. flexneri* induces robust LDH release from infected RAW264.7 macrophages, which is reduced in infections with a Δ*ipaH7.8* mutant (Fig. 4F). Cell killing by the Δ*ipaH7.8* strain was complemented with a plasmid expressing *ipaH7.8*. Using CRISPR/Cas9-engineered RAW264.7 cells (*47*), we found IpaH7.8-dependent cell death is markedly reduced in cells lacking CASP1 or NLRP1B. The NLRC4 inflammasome also recognizes *S. flexneri* (*48-50*). In *Nlrc4*^−/−^ RAW cells, inflammasome activation was almost entirely IpaH7.8-dependent (Fig. 4F). As expected, direct ubiquitylation of NLRP1B by *S. flexneri* circumvents the requirement for the N-end rule ubiquitin ligase *Ubr2* identified by the Bachovchin group to be required for LF-mediated NLRP1B activation (*Fig. S6*). Immortalized 129 macrophages were also sensitive to IpaH7.8-dependent killing, which correlated with decreased levels of endogenous NLRP1B and the induction of CASP1 maturation (Fig. 4G). *S. flexneri* is not a natural pathogen of mice; this may be due in part to species-specific NLRP1B effector recognition as human NLRP1 does not appear to detect IpaH7.8 (*Fig. S6D*). Taken together, our new mechanistic understanding of NLRP1B has led us to identify ubiquitin ligases as a new category of pathogen-encoded enzyme that activates NLRP1B.

Prior work in *Arabidopsis* has shown that an NLR called RPS5 detects proteolytic cleavage of the host PBS1 kinase by the translocated *Pseudomonas syringae* effector AvrPphB (*51*). In this system, RPS5 appears to detect the cleavage products of PBS1 (*52*). Although NLRP1B also appears to detect a pathogen-encoded protease, our results suggest that the underlying mechanism is very different than that of RPS5. Instead of proteolysis generating a specific ligand, it appears that NLRP1B is itself the target of proteolysis, leading to its proteasomal degradation, and release of a functional inflammasome fragment. Thus, NLRP1B is in essence a sensor of its own stability, permitting detection of diverse pathogen-encoded enzymes, potentially including those of viruses or parasites (*53-55*). Although IpaH7.8 and LF protease both activate NLRP1B, we favor a scenario in which the ‘intended’ targets of these pathogen-encoded enzymes are other host proteins, and that NLRP1B has evolved as a decoy target of these proteins. Although decoy sensors are widely deployed in plant immunity (*56*), NLRP1 may be the first such sensor described in animals.

NLRP1 was the first protein shown to form an inflammasome (*11*). Our results provide a long-sought mechanism that explains how NLRP1 is activated. Our proposed mechanism may also apply to other FIIND-death domain fold containing proteins, including PIDD1 and CARD8 (*19*). In addition to explaining how NLRP1 senses pathogens, our new mechanistic understanding also likely provides an explanation for why naturally occurring mutations that destabilize human NLRP1 (*27*) also result in its activation. It has previously been suggested that NLRP1B is activated upon sensing of ATP-depletion in cells (*38, 45*). Our results are not incongruous with this model; indeed, ATP-depletion may also indirectly affect NLRP1B stability. Taken together, our results lay the foundation for identifying pathogen-encoded activators of human NLRP1 and provide a conceptual basis for designing therapeutic interventions that target NLRP1.

## Acknowledgments

We are grateful to J. Chavarría-Smith for discussions and for laying the experimental foundations for this work. We thank P.R. Beatty and UC Berkeley undergraduates in the MCB150L course for help in generating the 2A12 monoclonal, the Rape Lab for guidance with ubiquitylation assays, the Bachovchin Lab for the HEK and RAW cell lines and for sharing results prior to submission, G. Barton, J. Chavarría-Smith, H. Darwin and J. Tenthorey for comments on the manuscript, and members of the Vance and Barton Labs for discussions.

## Author contributions

Conceptualization, A.S., P.S.M., R.E.V.; Methodology, A.S., P.S.M., R.E.V.; Investigation, A.S., P.S.M., L.G., E.W.M.; Resources, A.S., P.S.M., L.G., C.F.L., R.E.V; Writing – Original Draft Preparation, A.S., P.S.M., R.E.V.; Writing – Review & Editing, A.S., P.S.M., C.F.L., R.E.V.; Visualization, A.S., P.S.M.; Supervision, A.S., P.S.M., C.F.L., R.E.V.

## Data and materials availability

All data is available in the main text or the supplementary materials.

## Supplementary Materials

### Materials and Methods

#### Plasmids and constructs

The coding sequence of *Nlrp1b* allele 1: 129S1/SvimJ (129) *Nlrp1b* (DQ117584.1), allele 2: C57BL/6J (B6) *Nlrp1b* (BC141354) or allele 3: AKR/J (a gift from Jeremy Mogridge) (*21*) and variants thereof, were cloned into either pCMSCV-IRES-GFP or pAcSG2, resulting in N-terminal maltose binding protein (MBP) tag followed by a 3C protease cleavage site and a C-terminal HA or FLAG tag, respectively, or into pQCXIP resulting in N-terminal GFP and C-terminal HA tags. The Fla-NLRP1B hybrid was constructed by replacing the first 45 N-terminal amino acids of NLRP1B with residues 431-475 of *Legionella pneumophila* Flagellin (ANN95373) followed by the TEV cleavage sequence. CASP1, IL1B, TEV and LF producing constructs were described previously (*16*). AID (Addgene #80076) and TIR1 (Addgene #80073) producing constructs were a gift from Andrew Holland. The *Nlrp1b* coding sequence was subcloned in-frame with AID-GFP. GFP-fused IpaH producing plasmids were constructed as follows: the *ipaH* coding sequences from the *S. flexneri* 2a str. 2457T virulence plasmid were transferred using the Gateway^TM^ vector conversion system (ThermoFisher) from Gateway entry clones (*57*) into the SmaI restriction site of the Gateway-compatible destination vector pC1-eGFP (Clontech) via LR reactions. The *ipaH7.8* coding sequence was also subcloned into pQCXIP resulting in an N-terminal mCherry tag. For protein expression in *E. coli* the *ipaH* coding sequences were subcloned into pET28a with a C-terminal 6X-HIS tag. Mutations were engineered by overlapping PCR.

#### Cell culture

293T and RAW264.7 cells were grown in DMEM supplemented with 10% FBS, 100U/ml penicillin, 100mg/ml Streptomycin and 2mM L-glutamine. Primary bone-marrow derived macrophages (BMDMs) were cultured in RPMI supplemented with 5% FBS, 5% mCSF, 100U/ml penicillin, 100mg/ml Streptomycin and 2mM L-glutamine. BMDM immortalization was previously described (*58*).

#### Bacterial strains and infections

2457T *S. flexneri*-derived *ipaH* deletion strains were constructed using the λ red recombinase-mediated recombination system (*59*), as previously described (*60*). To construct complemented strains, the coding sequence of *ipaH7.8* and 407 basepairs upstream, representing the endogenous promoter, were Gateway cloned into the pCMD136 plasmid and transformed into the Δ*ipaH7.8* mutant strain. *S. flexneri* was grown at 37°C on tryptic soy agar plates containing 0.01% Congo red, supplemented with 100 μg/ml spectinomycin for growth of complemented strains. For infections, 5 ml tryptic soy broth (TSB) was inoculated with a single Congo red-positive colony and grown overnight shaking at 37°C. Saturated cultures were back-diluted 1:100 in 5 ml fresh TSB and incubated for 2-3 hours shaking at 37°C. Bacteria were washed in cell culture medium and spun onto cells for 10 minutes at 300×*g*. Infected cells were incubated at 37°C for 20 minutes and then washed twice with cell culture medium containing 25μg/ml gentamicin, then returned to 37°C for further incubation (30 minutes to 2 hours). Cells were infected at an MOI of 30 unless otherwise specified. Cell death was assessed by LDH activity in clarified culture supernatants as previously described (*61*). Protein in supernatants was TCA precipitated for anti-CASP1 immunoblotting.

#### Reconstituted NLRP1B activity assays

To reconstitute inflammasome activity in 293T cells, constructs producing NLRP1B (or mutants), CASP1 and IL-1β were co-transfected with constructs producing TEV, LF, IpaHs or empty vector (MSCV2.2 or pcDNA3) using Lipofectamine 2000 (Invitrogen) following the manufacturer’s protocol. For experiments using recombinant proteins, fresh media containing 10μg/ml PA and 2.5μg/ml LF, supplemented with or without 10μM MG132, 1μM Bortezomib, or 0.5μM NMS-873, was added to cells for 2-4 hours. For auxin-inducible degradation, AID-NLRP1B and TIR1-producing constructs were co-transfected and treated with 500μM indole-3- acetic acid sodium salt (IAA) (Sigma) for 3-6h in the presence or absence of 10μM MG132. In all experiments, cells were lysed in RIPA buffer with protease inhibitor cocktail (Roche) 24 hours post-transfection.

#### Endogenous NLRP1B activity assays

2.5×10^6^ i129 BMDMs were plated in a 6-well plate. 2 hours prior to challenge, cells were primed with 1.0μg/ml Pam3CSK4 (Invivogen). Cells were washed with PBS and media was replaced with 0.5ml Opti-MEM (Gibco) with or without 20μg/ml PA, 10μg/ml LF and/or 10μM MG132. Cells and media were lysed by addition of 120μl of 10X RIPA buffer with protease inhibitor cocktail 2.5 hours post-treatment.

#### Immunoblotting and antibodies

Lysates were clarified by spinning at ~16000×*g* for 10 minutes at 4°C. Clarified lysates were denatured in SDS loading buffer. Samples were separated on NuPAGE Bis-Tris 4-12% gradient gels (ThermoFisher) following the manufacturer’s protocol. Gels were transferred onto Immobilon-FL PVDF membranes at 35V for 90 minutes and blocked with Odyssey blocking buffer (Li-Cor). Proteins were detected on a Li-Cor Odyssey Blot Imager using the following primary and secondary antibodies: anti-HA clone 3F10 (Sigma), anti-IL-1β (R&D systems, AF-401-NA), anti-MBP (NEB, E8032S), anti-GFP (Clontech, JL8), anti-mCherry (ThermoFisher, 16D7), anti-CASP1 (Adipogen, AG-20B-0042-C100), anti-Ubiquitin (Cell Signaling, P4D1), Alexfluor-680 conjugated secondary antibodies (Invitrogen). Band intensities were quantified with Image Studio Lite software v5.2.5. The complete gel images are shown in *Fig. S7*.

To produce the 2A12 mouse anti-NLRP1B monoclonal antibody, the pAcSG2-*Nlrp1b* construct was co-transfected with BestBac linearized baculovirus DNA (Expression Systems) into SF9 cells following the manufacturer’s protocol to generate NLRP1B expressing baculovirus. Primary virus was amplified in SF9 cells. NLRP1B was produced by infecting 4L of High Five cells with 1ml of amplified virus/L cells. Cells were harvested 48 hours after infection by centrifugation at 300×*g* for 15 minutes. Cell pellets were resuspended in lysis buffer (50mM Hepes pH7.5, 150mM NaCl, 1% NP-40, 5% glycerol) and lysed on ice using a dounce homogenizer. Homogenized samples were clarified at 24000x*g* for 30 minutes and supernatants were batch bound to 1ml amylose resin for 2 hours at 4°C. Samples were column purified by gravity. Resin was washed with 50ml of wash buffer (20mM Hepes pH7.4, 150mM NaCl, 0.02% NP-50, 5% glycerol). Sample was eluted with 1ml fractions of elution buffer (20mM Hepes pH7.4, 150mM NaCl, 0.02% NP-50, 5% glycerol, 20mM Maltose). Peak elutions were pooled and MBP was cleaved by treatment overnight with 3C protease. Free MBP was removed by passing the sample over amylose resin. BALB/c mice were immunized with 10μg NLRP1B plus 100μl Sigma adjuvant four times. Splenocytes were fused the with the P3X63-Ag8.653 parental line. Clones were screened via ELISA against recombinant NLRP1B protein or recombinant FLAG-tagged MBP protein to identify clones specifically reactive to NLRP1B. Clarified supernatant from the hybridoma clone 2A12 was used for immunoblotting.

#### In Vitro Ubiquitylation Assay

Recombinant 129 or B6 NLRP1B was produced in insect cells and purified as described above prior to 3C treatment. Recombinant IpaH7.8, IpaH7.8C357A (catalytic mutant), and IpaH9.8 were expressed in BL21 *E. Coli*. BL21. 1L of cells were grown to ~0.7 OD600 and induced with 1M IPTG (Sigma) for 4 hours at 37 °C. Pellets were resuspended in 50mM Tris pH7.4, 150mM NaCl, 1%NP-40 and sonicated to lyse. Samples were clarified at 24000×*g* for 30min. NaCl concentration was of the supernatants was increased to 400mM and 20mM imidazole pH8.0 was added to samples. Supernatants were batch bound to 1ml Ni resin (Qiagen) at 4°C for 2 hours. Samples were purified by gravity, washed with 50ml of 20mM Tris pH7.4, 400mM NaCl, 20mM imidazole pH8.0. Protein was eluted in 1ml fractions of 20mM Tris pH7.4, 150mM NaCl, 250mM imidazole pH8.0. Elution peaks were pooled and desalted into 20mM Hepes pH7.4, 150mM NaCl, 2mM DTT.

In vitro ubiquitylation assays were performed in 25mM Tris pH7.4, 50mM NaCl, 10mM MgCl_2_, 5mM ATP, 0.1mM DTT with 60nM Ubiquitin E1 (Boston Biochemistry), 200nM UbcH5c (Boston Biochemistry), 10μM Ubiquitin (Boston Biochemistry), 300nM IpaH, and 270nM NLRP1B. Reaction was run for 1 hour at 37 °C. Solutions were then batch bound to anti-FLAG M2 agarose gel (Sigma) at 4 °C for 2 hours. Bound samples were column purified, washed with 5ml of HBS (20mM Hepes, 150mM NaCl). Final samples were eluted in 150μl HBS+150μg/ml FLAG peptide.

#### Native gel oligomerization assay

Samples were transfected into 293T cells with constructs as above in a 6-well plate. After 24 hours, samples were harvested by removing media and washing cells off plate with cold PBS. Harvested cells were centrifuged at 300x*g* for 10 minutes at 4°C. Cells were lysed in lysis buffer (50mM Hepes pH7.5, 150mM NaCl, 1% NP-40, 5% glycerol) and samples were clarified by spinning at 16000x*g* for 10 minutes at 4°C. Samples were run on NativePAGE Bis-Tris gels (ThermoFisher) according to manufacturer’s protocols.

#### Detection of UPA-CARD and ASC speck formation by immunofluorescence

293T cells were grown on fibronectin-coated coverslips. Constructs producing NLRP1B and ASC were co-transfected with constructs producing TEV or an empty vector. The NLRP1B construct was designed with a C-terminal FLAG and N-terminal HA tag, where the HA sequence was inserted following the P1′ position of the TEV cleavage site. 20h post-transfection, cells were fixed with 4% PFA / PBS (20min) and permeabilized in 0.5% saponin / PBS (5min). Blocking and antibody staining was performed at room temperature in 5% BSA / 0.1% saponin /PBS. Primary antibodies: rabbit anti-ASC (Santa Cruz, N-15), rat anti-HA (Roche, 3F10), mouse anti-FLAG (Sigma, M2). Secondary antibodies: goat anti-rabbit AMCA (Jackson Laboratories), goat anti-rat Alexa Fluor 647, goat anti-mouse Alexa Fluor 555 (Molecular Probes). Coverslips were mounted onto slides using Vectashield medium (Vector Laboratories, Inc., H-1000), and imaged with a ZEISS LSM 710. To quantify the presence of FLAG or HA per ASC speck, the Imaris imaging software (Bitplane) was used to first identify ASC specks as objects, then to count FLAG- and/or HA-positive objects based on a fluorescence intensity threshold set manually for each channel. Thresholds were determined manually using a training set of samples and controls (i.e., no primary antibody), and applied in batch to all samples.

---

**Fig. S1.**
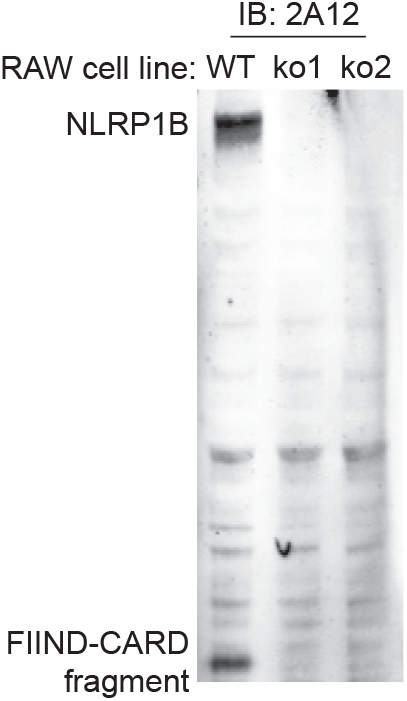
(Fig. 2) The 2A12 monoclonal antibody recognizes the CARD domain of NLRP1B. NLRP1B was detected using the anti-NLRP1B monoclonal Ab 2A12 in lysates from WT or *Nlrp1b^−/−^* RAW264.7 cells. ko1 and ko2, CRISPR-Cas9 *Nlrp1b^−/−^* cells clone 1 and 2 (*47*).

**Fig. S2.**
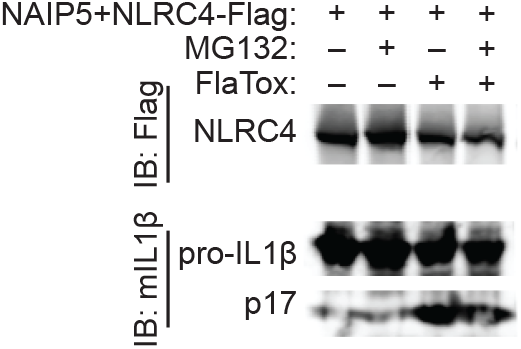
(Fig. 2) Proteasome inhibitors do not block NAIP/NLRC4 inflammasome activation. The proteasome is specifically required for NLRP1B inflammasome activation. The proteasome inhibitor MG132 (10μM) does not block ligand-induced NAIP-NLRC4 inflammasome activation in reconstituted 293T cells. FlaTox, PA + LFn-FlaA (*62*).

**Fig. S3.**
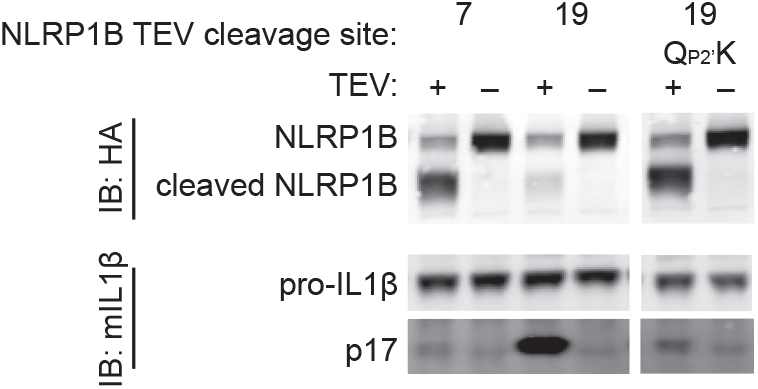
(Fig. 1) P2′ residue identity modulates TEV-induced proteasomal degradation and activation of NLRP1B. TEV cleavage at site 7 and site 19 differentially induce degradation and activation of NLRP1B, despite having the same neo-N-terminal amino acid following TEV cleavage. To determine whether the local amino acid environment effects NLRP1B stability and activation following TEV cleavage, the P2′ position from the relatively inactive (site 7, P2′=K) was introduced into the active (site 19, P2′=Q) cleavage site. The activity of the indicated C-terminal HA-tagged variants was tested in 293T cells as described in Fig. 1. A P2′ lysine is sufficient to both stabilize and inactivate NLRP1B at site 19, suggesting that other residues beyond the P1′ position is important for determining the stability (and subsequent activation) of the NLRP1B N-terminus.

**Fig. S4.**
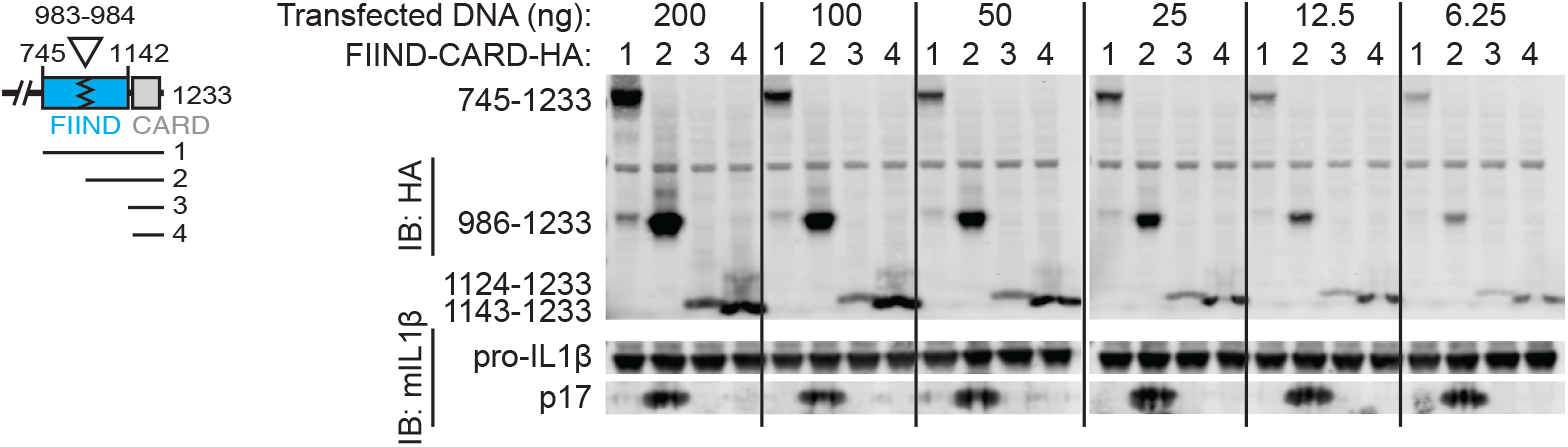
(Fig. 3) The FIIND(UPA)-CARD fragment is a potent activator of IL1β processing. The activity of C-terminal HA-tagged variants corresponding to the full length FIIND(ZU5+UPA)-CARD [(1), 745-1233), FIIND(UPA)-CARD fragment [(2), 986-1233), FIIND(UPA)-CARD truncation [(3), 1124-1233) or the CARD domain [(4), 1143-1233) were transfected across a range of ng inputs in 293T cells, as described for Fig. 3B. Only the FIIND(UPA)-CARD fragment exhibited activity.

**Figure S5.**
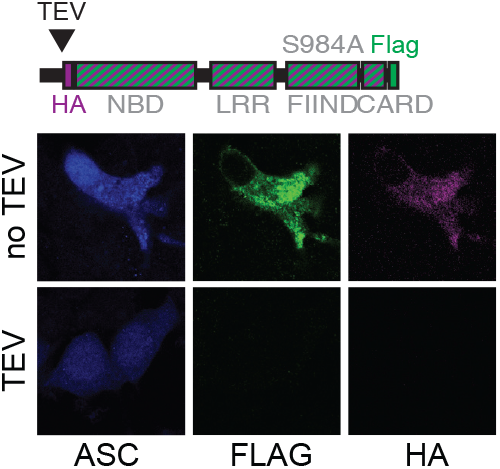
(Fig. 3). FIIND auto-processing is required for release of the FIIND(UPA)-CARD and its colocalization to the ASC speck. 293T cells were transfected with constructs producing ASC (blue) and an NLRP1B FIIND mutant (S984A) variant marked with a C-terminal FLAG (green) and an N-terminal HA (magenta). As schematized, the HA is inserted directly after the TEV cleavage site, allowing detection of the TEV-cleaved protein. Representative images depict cytosolic FLAG and HA signal in untreated and TEV-expressing cells. Unlike in Fig. 3G, where ASC colocalizes specifically with FLAG and not HA, there was no specific FLAG-ASC staining was observed for the FIIND mutant protein.

**Figure S6.**
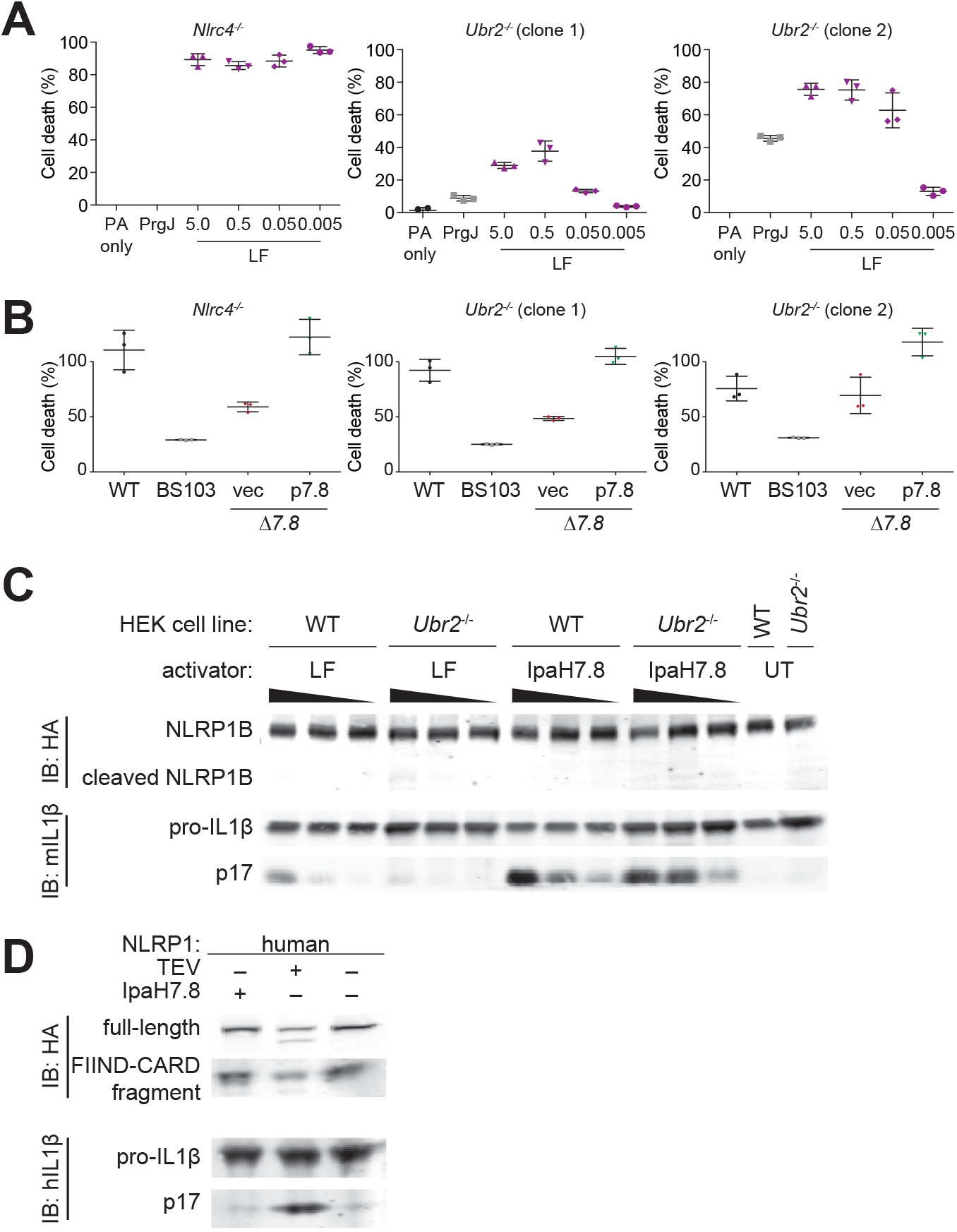
(Fig. 4) IpaH7.8 activates mouse NLRP1B, but not human NLRP1, independently of N-end rule ubiquitin ligases. **(A)** LF but not IpaH7.8-mediated macrophage cell death requires UBR2. PrgJ or varying amounts of LF (μg/mL) was delivered into *Nlrc4^−/−^* or *Ubr2^−/−^* RAW264.7 cells. LF-mediated cell death remains saturated in *Nlrc4^−/−^* cells at all LF concentrations, but is reduced in two *Ubr2^−/−^* cell lines at lower concentrations. (**B**) IpaH7.8-mediated cell death is independent of UBR2. Infection (MOI 10) of *Nlrc4^−/−^* or *Ubr2^−/−^* RAW264.7 cells with the WT *Shigella flexneri* strain 2457T (black circle) or mutants strains. BS103, virulence plasmid-cured (grey box); vec, Δ*7.8* strain complemented with pCMD136 empty vector (red diamond); Δ*7.8* strain complemented with pCMD136 *ipaH7.8* (green inverted triangle). Cell death was monitored by assaying for lactate dehydrogenase (LDH) activity in culture supernatants (A and B). (**C**) LF, but not IpaH7.8, activation of the NLRP1B inflammasome requires UBR2. The NLRP1B inflammasome was reconstituted in 293T cells as described in Fig. 1. CASP1-dependent processing of IL1β to p17 induced by LF, but not IpaH7.8, was reduced in *Ubr2^−/−^* cells relative to WT cells. UT, untransfected. (**D**) IpaH7.8 does not degrade or activate human NLRP1. The human NLRP1inflammasome was reconstituted in 293T cells, and inflammasome activation was monitored by human CASP1 processing of human pro-IL-1β to p17. A TEV-cleavable human NLRP1 variant was used to allow for TEV-mediated inflammasome activation.

**Figure S7.**
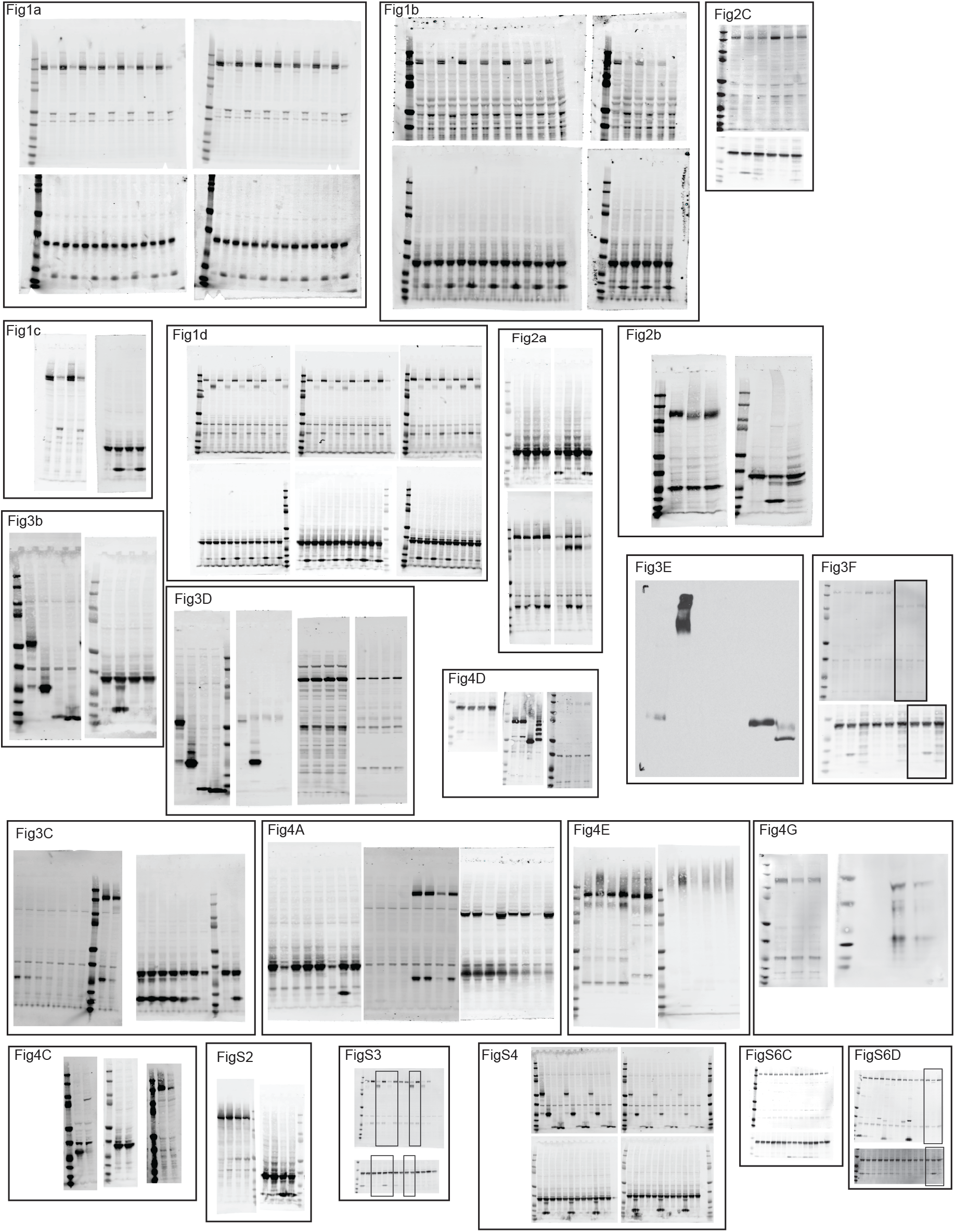
Complete images for all gels. Above are shown the full images for each gel presented.

